# Correlating enzyme annotations with a large set of microbial growth temperatures reveals metabolic adaptations to growth at diverse temperatures

**DOI:** 10.1101/271569

**Authors:** Martin KM Engqvist

## Abstract

Interpreting genomic data to identify temperature adaptations is challenging due to limited accessibility of growth temperature data. In this work I mine public culture collection websites to obtain growth temperature data for 21,498 organisms. Leveraging this unique dataset I identify 319 enzyme activities that either increase or decrease in abundance with temperature. This is a striking result showing that up to 9% of enzyme activities may represent metabolic changes important for adapting to growth at differing temperatures in microbes. Eight metabolic pathways were statistically enriched for these enzyme activities, further highlighting specific areas of metabolism that may be particularly important for such adaptations. Furthermore, I establish a correlation between 33 domains of unknown function (DUFs) with growth temperature in microbes, four of which (DUF438, DUF1524, DUF1957 and DUF3458_C) were significant in both archaea and bacteria. These DUFs may represent novel, as yet undiscovered, functions relating to temperature adaptation.

## Introduction

The physiology and evolutionary adaptations of living organisms are shaped by the environmental factors that define the habitat in which they live, but availability of metadata describing the environment of each habitat is limited. This is true for such properties as growth temperature, pH, salinity and many more. Several projects addressing aspects of this limitation have been published in recent years. The most comprehensive of these is BacDive (Söhngen et al. 2014, 2015) which was created by digitizing and mining analog records containing microbial metadata (Reimer et al. 2017). Other resources include MediaDB, which offers a hand-curated database for microbial growth conditions (Richards et al. 2014). Records regarding the phenotypic and environmental tolerance for over 5,000 species has also been made available (Barberán et al. 2017). One recent approach leveraged text mining to infer a large number of phenotypic traits from the scientific literature and the World Wide Web (Brbić et al. 2016).

Among environmental factors temperature is unique in that it crosses physical barriers. As a result, organisms cannot efficiently shield themselves from temperature in the way they can shield themselves from extreme external pH or salinity by maintaining steep concentration gradients over biological membranes. Instead, each biomolecule inside microorganisms must be adapted to the temperatures in which they grow. This makes proteins and metabolites from microbes growing at hot temperatures particularly interesting for biotechnological and industrial applications (Mehta et al. 2016). Organisms can be generally categorized based on the temperature range of their optimal growth. No consensus has been reached regarding the exact temperature range of each category, here I make use of the following: psychrophiles (< 15 °C), mesophiles (15-50 °C), thermophiles (50-80 °C) and hyperthermophiles (> 80 °C).

A range of specific adaptations to high temperatures are known, with many studies focusing on characteristics of thermophilic genomes or molecules such as proteins and lipids (Wang, Cen, and Zhao 2015; Stetter 1996). Stability of thermophilic proteins is attributed to their hydrophobic cores, increased numbers of charged residues and disulphide bonds (Wang, Cen, and Zhao 2015; Boutz et al. 2007; Bezsudnova et al. 2012). Cell membranes in thermophiles are characterised by high permeability barrier and capacity to maintain the liquid crystalline phase, due to presence of saturated fatty acids in bacteria, and ether lipids in archaea (Koga 2012). Adaptations on the DNA level are also known. For example, genomes of thermophiles are generally smaller than those of mesophiles, with reduced number of some protein family members (van Noort et al. 2013; Burra, Kalmar, and Tompa 2010; Sabath et al. 2013). Horizontal gene transfer is thought to be important driving force for thermophilic adaptation. For instance, reverse gyrase, which was shown to have heat-protective DNA chaperone activity (Kampmann and Stock 2004), is considered to be transferred from archaea to bacteria (Forterre et al. 2000; Aravind et al. 1998).

Metabolism varies widely among thermophiles and general trends are hard to discern as it is extremely difficult to distinguish between the effect of speciation versus adjustment to extreme environments. For example, three main glycolytic pathways are used by bacteria in general: the traditional Embden-Meyerhof (EM) glycolysis, the Entner-Doudoroff pathway, and the pentose phosphate pathway; all three can also be found in different thermophilic bacteria (Counts et al. 2017; Swarup et al. 2014; Brumm et al. 2015; Selig et al. 1997). In archaea, however, only modified variants of classical sugar degradation pathways were identified (Brasen et al. 2014; Selig et al. 1997). For example, *Pyrococcus furiosus* contains a nontraditional variation of EM glycolysis, in which ADP-dependent kinases are involved and glyceraldehyde-3-phosphate is converted directly to 3-phosphoglycerate, with no thermolabile 1,3-biphosphoglycerate intermediate present as in traditional glycolysis (Kengen et al. 1994; Ettema et al. 2008). Even within the same taxonomic domain, sets of used pathways may vary. Flux analysis of three extremely thermophilic bacteria indicated that the metabolisms of the studied thermophiles were highly distinct, with differences in such pathways as amino acid or NADPH metabolism (Cordova et al. 2017). However, a communality is that all three strains relied heavily on glycolysis and the TCA cycle. Studies on *Thermus thermophilus,* possibly the most well-studied thermophile, revealed alternative pathways of amino acids synthesis, with lysine being synthesized by alpha-aminoadipate pathway instead of diaminopimelate pathway (Kosuge and Hoshino 1998) and homocysteine produced from an alternative precursor, O-acetyl-L-homoserine (Lee et al. 2014). *T. thermophilus* is also known to produce a variety of polyamines, the most common ones, spermidine and spermine, are synthesised using a distinct pathway from L-arginine via aminopropyl agmatine (Oshima 2007). Analysis of complete thermophilic genomes is a widely used method of finding both novel metabolic pathway, as well as enzymes of potential biotechnological use (Henne et al. 2004; Schäfers et al. 2017; Wu et al. 2009). An alternative pathway of menaquinone (MK) synthesis was discovered when no genes from the classical pathway were found in genomes of some MK-producing microorganism (Hiratsuka et al. 2008). Genes involved in the novel MK synthesis pathway are present in a range of thermophilic microorganisms, which may play a role in adaptation to growth at high temperatures, as one of the intermediates in the classical pathway, isochorismate, is known to be thermolabile (Fang, Langman, and Palmer 2010).

Several studies avoid the limitations associated with comparing a small number of thermophilic versus mesophilic species, thereby identifying more general trends, by performing metagenomic analysis of environmental samples from extreme habitats (DeCastro, Rodríguez-Belmonte, and González-Siso 2016; Cowan et al. 2015). A study comparing metagenomes obtained from cold and hot deserts found thermophiles having a higher number of genes involved in metabolism and transport of carbohydrates and secondary metabolites (Le et al. 2016). Similar results were found in other studies of metagenomes from hot springs in India, where abundance of KEGG (Ogata et al. 1999; Kanehisa et al. 2017) pathways was investigated (Badhai, Ghosh, and Das 2015; Saxena et al. 2016).

In this work I investigate metabolic trends in adapting to thermophilicity by comparing the presence of metabolic enzymes over many thousands of organisms, growing at various temperatures. I identify 319 individual enzymatic reactions that are preferentially present at either low or high temperatures, indicating metabolic adaptations. We identify eight pathways which are over-represented in metabolic changes. These may represent parts of metabolism that are particularly important for the adaptation to growth at different temperatures. Finally, I show that 33 protein domains of unknown function (DUFs) likewise correlate with growth temperature, a result that may provide important clues to their function. Enabling this approach is a unique dataset of 21,498 organisms with their growth temperatures, which I obtained through data mining of publicly available organism culturing protocols from major culture collection centers. I make this data available to enable researchers to gain additional insights into evolutionary temperature-dependent adaptations.

## Results

### Culture collection center websites are a rich source for growth temperature data

The hypothesis underlying this project is that, given a sufficiently large dataset with organism growth temperatures, meaningful correlations between growth temperatures and biological adaptations can be made. I reasoned that publicly available culturing protocols from microorganisms in major stock collection centers may form a resource that could be mined for such growth temperatures. Custom software in Python (http://www.python.org) was designed to systematically download all of the public web pages containing organism information from four stock collection centers: ATCC (http://www.lgcstandards-atcc.org), DSMZ (http://www.dsmz.de), NCTC (http://www.phe-culturecollections.org.uk) and NIES (http://www.shigen.nig.ac.jp). Each of these pages were then mined for the organism name and the growth temperature (Figure 1A). A fifth stock center, the Institute Pasteur stock collection (http://www.research.pasteur.fr/en/team/biological-resources-center/) provided an excel table with organism names and growth temperatures upon request. Finally, organism growth temperatures from the BacDive database (Söhngen et al. 2014, 2015) were collected using scripts interfacing with the public API.

**Figure 1.**
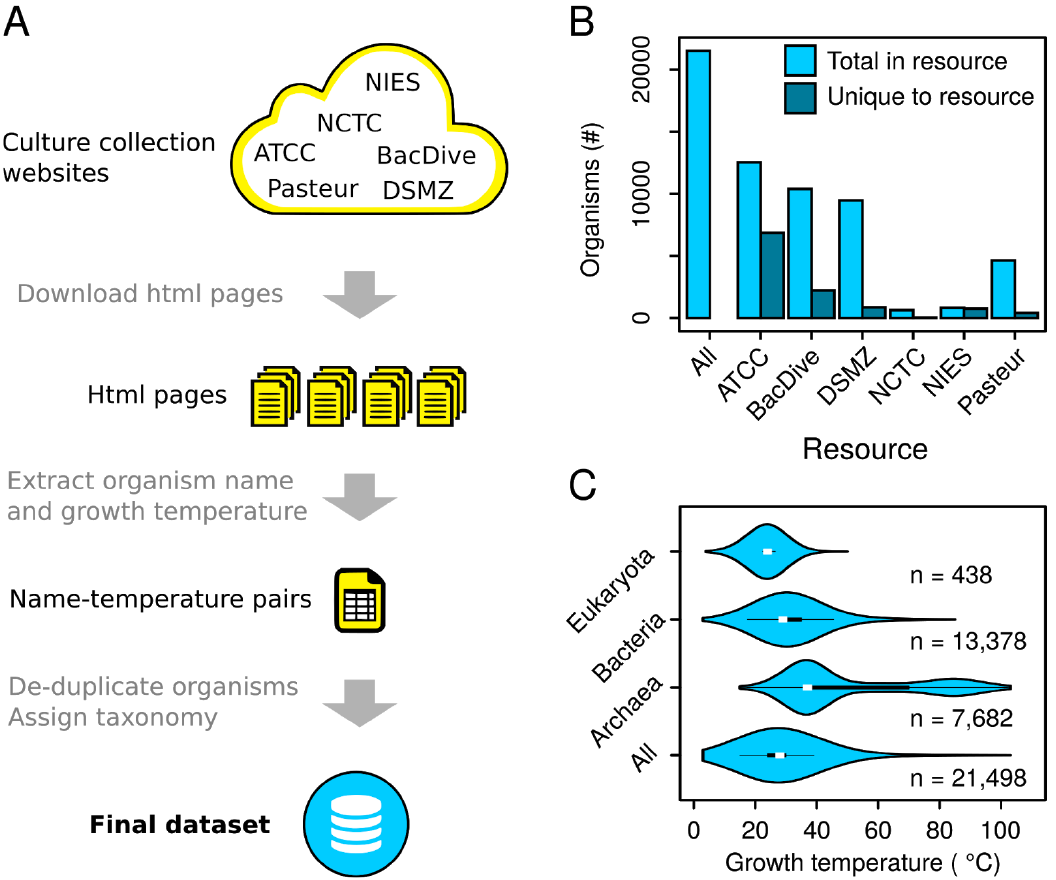
Culture collection centers represents a rich source for growth temperature data. **(A)** An overview schematic showing how the growth temperature dataset was obtained. **(B)** A database comparison showing which provided the largest number of unique organisms not present in the others. Total in resource shows the total number of organisms in each database, unique to resource shows how many organisms in that database were unique. **(C)** A violin-plot showing the growth distribution of growth temperatures in the dataset for each of the three domains of life. The white dot represents the median. The broad black bar represents the upper and lower quartile, which contain 50% of the data points. The thin black lines represent the upper and lower adjacent values. The outer plot shape is a kernel density plot that visualizes the probability distribution of the data.

Organism names were collapsed to the species level (ignoring strain designations) and the number of organisms in each of the databases was compared (Figure 1B). A total of 21,498 unique organism names combined with their growth temperature were obtained. ATCC and BacDive had the greatest number of unique organisms that were not present in any other database, with 6,852 and 2,211 respectively. In order, DSMZ, NCTC, NIES and Pasteur had 854, 44, 749 and 415 organisms that were uniquely present in each of the databases.

Each organism name in the dataset was mapped to a taxonomic identifier using the NBCI taxonomy resource (https://www.ncbi.nlm.nih.gov/taxonomy). The data from all databases were combined into one non-redundant dataset, which is referred to as the growth temperature dataset. The majority of organisms in the dataset (62%) is made up of bacteria (Figure 1C). Most of these are psychrophiles and mesophiles. Only a small number are thermophiles. Archaea make up 36% of all organisms in the dataset, with a more even distribution between mesophiles, thermophiles and hyperthermophiles. Eukaryotes, mainly comprising fungi and protists, make up only 2% of the dataset and encompass both psychrophiles and mesophiles. In general the dataset contains many more psychrophiles and mesophiles than thermophiles and hyperthermophiles (Figure 1C). The growth temperature dataset is made available for re-use by other researchers as a tab-delimited file as well as in the xml format on Zenodo (https://zenodo.org/; doi: 10.5281/zenodo.1175609).

### Growth temperatures correlate strongly with mean enzyme optima

The growth temperature dataset was validated by investigating the correlation between the growth temperature of each organism and the temperature optima of enzymes from that organism. For this validation all experimentally determined enzyme temperature optima were extracted from the BRENDA enzyme database (Schomburg et al. 2017), https://www.brenda-enzymes.org). The resulting data contained experimentally determined temperature optima for 31,826 enzymes, sourced from 3,421 organisms. This dataset is referred to as the enzyme temperature dataset. The overall distribution of growth temperatures (Figure 2A) differ compared to the overall distribution of enzyme temperature optima (Figure 2B). The majority (88%) of growth temperatures fall in the range between 20 °C and 40 °C, with an abrupt drop in the number of organisms grown at temperatures over 40 °C. In comparison, the majority of enzyme temperature optima also fall in the range between 20 °C and 40 °C (64%), but their distribution decreases much less drastically above this temperature range.

**Figure 2.**
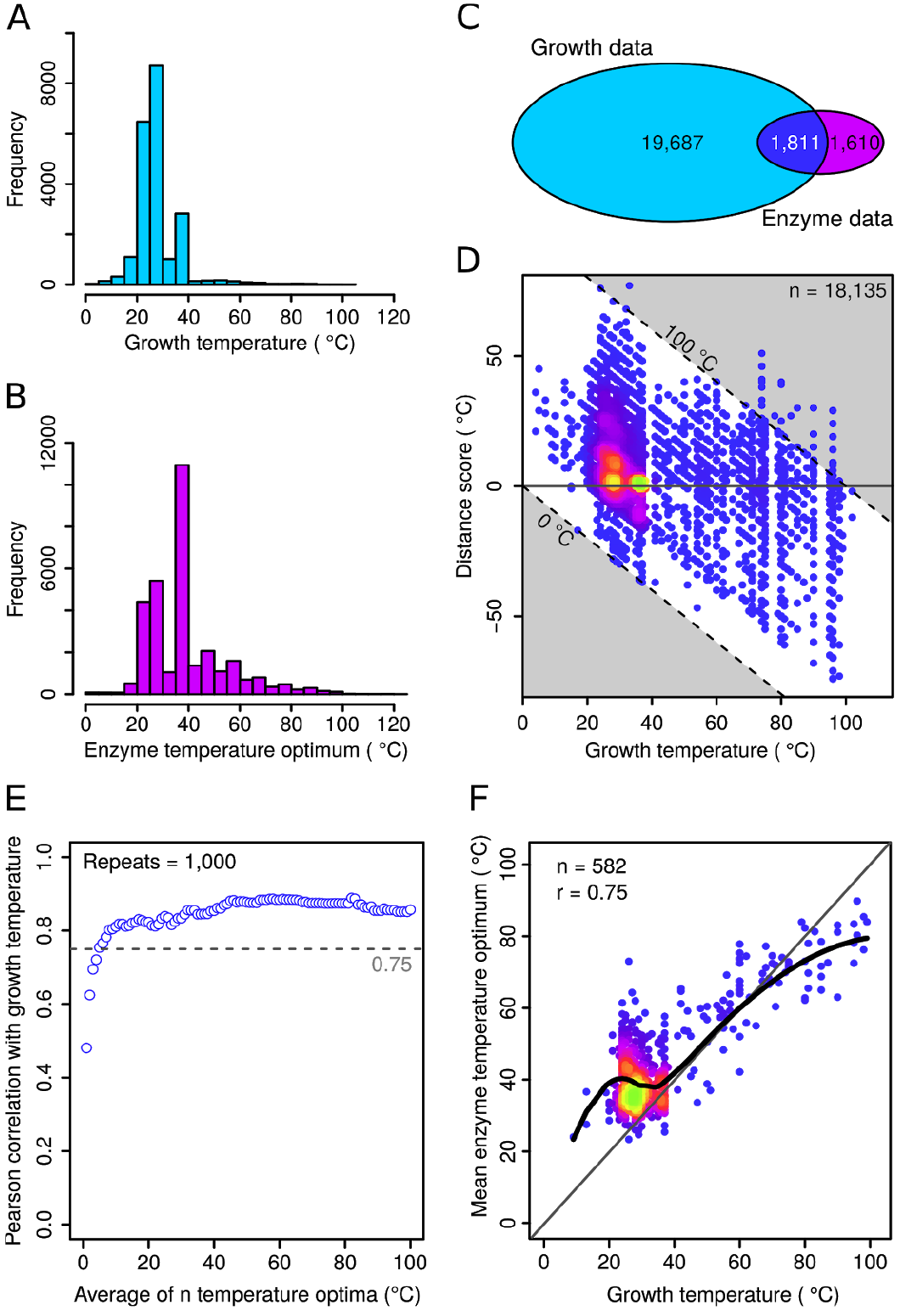
Growth temperatures correlate strongly with mean enzyme optima. **(A)** The distribution of growth temperatures for all organisms in the growth temperature dataset. **(B)** The distribution of experimentally determined optima for the 31,826 enzymes obtained from the BRENDA database. **(C)** A Venn diagram showing the number of organisms present in each of the growth temperature dataset, the enzyme temperature dataset, as well as the number of organisms present in both. **(D)** Distance score of individual enzyme temperature optima with growth temperature. The horizontal gray line indicates perfect agreement between the two values. The two diagonal dashed lines indicate absolute enzyme temperature optima of 0 °C and 100 °C. The colors indicate the spot density with brighter and more green colors indicating higher density. **(E)** A sensitivity plot showing the Pearson correlation coefficient between enzyme temperature optima and growth temperatures. Each data point represents the correlation coefficient obtained between the mean temperature optimum of n sampled enzymes in the range 1 <= n <= 100 and growth temperature. The average of five enzymes or more achieves a correlation coefficient above 0.75. **(F)** A scatterplot comparing the growth temperature of organisms organisms with more than five reported enzyme temperature optima with the mean temperature optimum of all enzymes reported for that organism. The diagonal gray line indicates perfect positive correlation. The thick black line represents a locally weighted polynomial regression. The Pearson correlation coefficient is given.

To be able to correlate growth temperatures with enzyme optima, organisms present in both datasets were identified (Figure 2C). Of the 21,498 organisms with growth temperatures and the 3,421 organisms with enzyme temperature optima only 1,811 were present in both datasets. From these the experimentally determined temperature optima for 18,135 enzymes were available. A simple distance score was computed by subtracting each organisms growth temperature from each individual enzymes temperature optimum (Figure 2D). Enzymes with optima exactly at the growth temperature have a distance score of 0 °C. Those which have an optimum higher than the growth temperature have a positive distance score, with a magnitude equal to the temperature difference. Enzymes with an optimum lower than the growth temperature have a negative distance score. Enzymes with optima between 20-100 °C could be found throughout the entire growth temperature range, indicating that not all enzymes match the organism growth temperatures (Figure 2D). Despite these deviations from the expected, half (51%) of the enzyme measurements were in fact within ± 10 °C of the growth temperature and 67% were within ± 15 °C of the growth temperature, showing that growth temperature may be a good indicator for the majority of enzyme optima.

This correlation was further investigated by comparing each organisms growth temperature with average enzyme optima calculated with an increasing number of averaged enzymes from that organism, ranging from 1 to 100 (Figure 2E). There is a clear trend for stronger correlation between growth and enzyme temperatures when increasing the number of averaged enzymes. This result shows that a single randomly chosen enzyme is an imprecise predictor of growth temperature (Figure 2E, bottommost circle). However, the mean optimum of at least five enzymes does display a Pearson correlation coefficient greater than 0.75. It is expected that the optimal catalytic temperature of enzymes follows growth temperature, the high correlation seen between these two variables is therefore a validation of the growth temperature dataset.

### Many enzyme activities correlate with temperature

A range of biological questions might be investigated by correlating biological properties with the collected organism growth temperatures. These could include genomic, transcriptomic, proteomic, metabolomic, phenotypic or taxonomic properties. To test this idea, I correlated enzyme activities, classified by Enzyme Commission numbers (EC numbers), with growth temperatures in an effort to identify temperature-dependent metabolic adaptations. A limitation in this approach is that it only focuses on known enzyme activities and novel ones cannot be directly identified. Missing annotations as well as mis-annotations also introduce noise in this analysis. Furthermore, it is important to consider that the presence of an enzyme coding sequence in a dataset does not mean it is expressed and functional in a given organism.

To obtain a set of enzyme activities for analysis all 88.6 million protein records from the UniProt database (UniProt Consortium 2015; https://www.uniprot.org/) were downloaded. Of these, 43% (38.3 million) could be annotated with the growth temperature of the organism from which they came - using the growth temperature dataset. EC numbers, were subsequently obtained for each of the matched records, where present, using the UniProt ID mapping tool (https://www.uniprot.org/uploadlists/). This resulted in the mapping of 3,551 unique EC numbers, which is approximately half of those currently listed in the BRENDA database. For the subsequent analysis Eukaryotes were excluded due to their limited range of growth temperatures in the dataset.

The growth temperature dataset is highly skewed toward organisms growing between 20 °C and 40 °C (Figure 1C). This skewing of the data would interfere with a correlation analysis. Therefore, for each individual EC number, the ratio of organisms having at least one protein carrying that annotation - versus those organisms that do not - was calculated to obtain a single value at each growth temperature. This results in a distribution of ratios over the analyzed temperatures for each individual EC number. A cutoff was set, including in the calculations only organisms with at least 1,000 protein records, as determined by a sensitivity plot (Figure 3A). This cutoff was used to remove noise caused by organisms with very few entries in UniProt. The proportion of proteins annotated as enzymes remained approximately constant across growth temperatures in both archaea and bacteria (Figure 3B).

**Figure 3.**
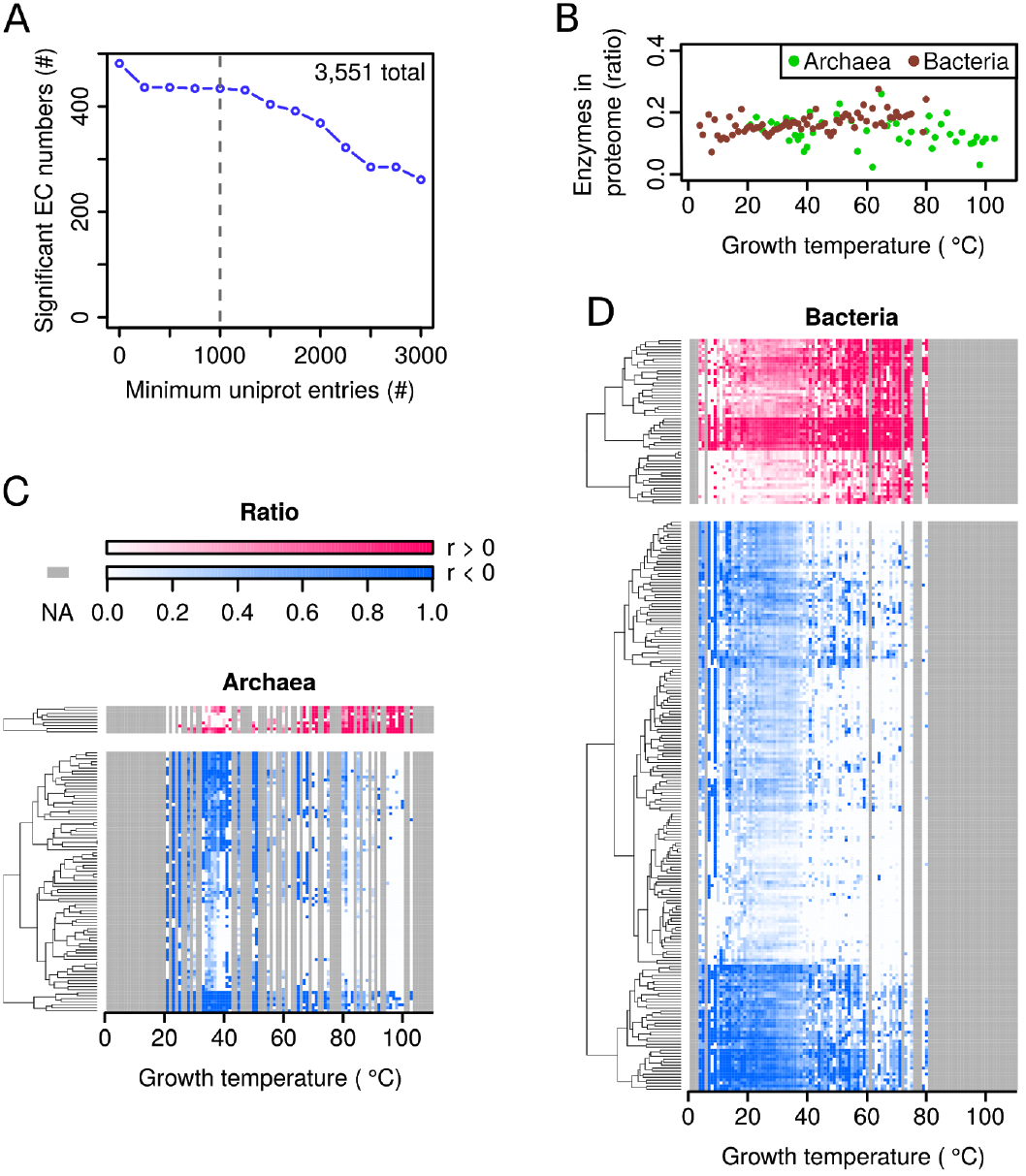
Many enzyme activities correlate with temperature. **(A)** A sensitivity plot showing the number of significant EC numbers (corrected p-value < 0.01) obtained at different cutoffs for minimum number of UniProt entries per organism. The gray dashed line indicates the selected cutoff. **(B)** Enzyme annotations as a proportion of the proteome. Each point represents the mean for all organisms growing a given temperature. The means for archaea and bacteria were calculated separately. **(C)** Heat-maps showing how the occurrence of significant EC numbers change with growth temperatures in archaea. Each row represents a different EC number and each column a different growth temperature. All included EC numbers are statistically correlated (corrected p-value < 0.01) either positively (red) or negatively (blue) with growth temperature and a minimum phylogenetic distance score of six (see Methods for details). For each reported growth temperature, the ratio of organisms containing each specific EC annotation is shown by the intensity of the color. Gray indicates temperatures for which no growth temperatures are available in the dataset. **(D)** Analysis and color scale as in C, but performed with data from bacteria.

In the growth temperature dataset there are proportionally more bacteria at low growth temperatures and proportionally more archaea at high growth temperatures (Figure S1). Correlating EC numbers and growth temperatures for the entire dataset may therefore highlight those that differ between these two domains of life, instead of those representing a true signal for temperature adaptation. bacteria and archaea were therefore analyzed separately. The Spearman correlation coefficient between the growth temperature and the enzyme occurrence ratios was calculated for each EC number and corrected p-values obtained (Figure S2). A concern in this analysis is that it may result in false positives representing enzyme activities only present in closely related organisms with a limited growth temperature distribution. To remove such false positives - and retain only those that may represent a general strategy of temperature adaptation - a filtering step based on phylogenetic distance was performed. Enzyme activities with significant correlation (corrected p-value below 0.01) were thus filtered to retain only those with a wide phylogenetic distribution, indicated by a phylogenetic distance score of at least six (see Methods for details).

From the 3,551 mapped EC numbers a total of 319 unique EC numbers remaining after filtering (340 total instances when accounting for recurring EC numbers), nine show statistically significant positive correlation in archaea and 55 in bacteria (Figure 3C and D, top panels; Supplemental file S1). Conversely, 86 enzymes from archaea show negative correlation with temperature and 190 in bacteria (Figure 3C and D, bottom panel; Supplemental file S1). 14 EC numbers recur in both the archaeal and bacterial dataset with the same correlation and 7 EC numbers show opposite correlation in archaea compared to bacteria. Together, these 319 EC numbers may shed light on metabolic adaptations important for growth at different temperatures.

### Certain metabolic pathways are enriched for the correlated enzyme activities

To take the analysis one step further I investigated whether any part of metabolism was enriched for the enzyme activities identified in the previous step. Such enrichment would implicate specific metabolic pathways, or sets of related pathways, in the process of adapting to differing growth temperatures. The enzyme activities were mapped onto KEGG (http://www.genome.jp/kegg) pathways and a hypergeometric test was applied to test for enrichment. Eight out of 155 KEGG pathways were statistically enriched for the enzyme activities. These were: TCA cycle (KEGG pathway map00020), ubiquinone and other terpenoid-quinone biosynthesis (KEGG pathway map00130), purine metabolism (KEGG pathway map00230), cysteine and methionine metabolism (KEGG pathway map00270), pyruvate metabolism (KEGG pathway map00620), one carbon pool by folate (KEGG pathway map00670), methane metabolism (KEGG pathway map00680), carbon fixation pathways in prokaryotes (KEGG pathway map00720). To highlight one of these pathways a portion of the “ubiquinone and other terpenoid-quinone biosynthesis pathway” is presented in Figure 4A with data from bacteria. The ubiquinone synthesis pathway contains three enzymes that show negative correlation with growth temperature (Figure 4B). Two different metabolic pathways lead to synthesis of menaquinone from chorismate. The first, so-called futalosine pathway, contains three enzymes that show positive correlation with temperature (Figure 4C). The second pathway of menaquinone biosynthesis contains five enzymes that show negative correlation with growth temperature and are not present at high temperatures (Figure 4D). These data suggest that an evolutionary adaptation to growth at high temperatures in many bacteria is to biosynthesize menaquinone via the futalosine pathway.

**Figure 4.**
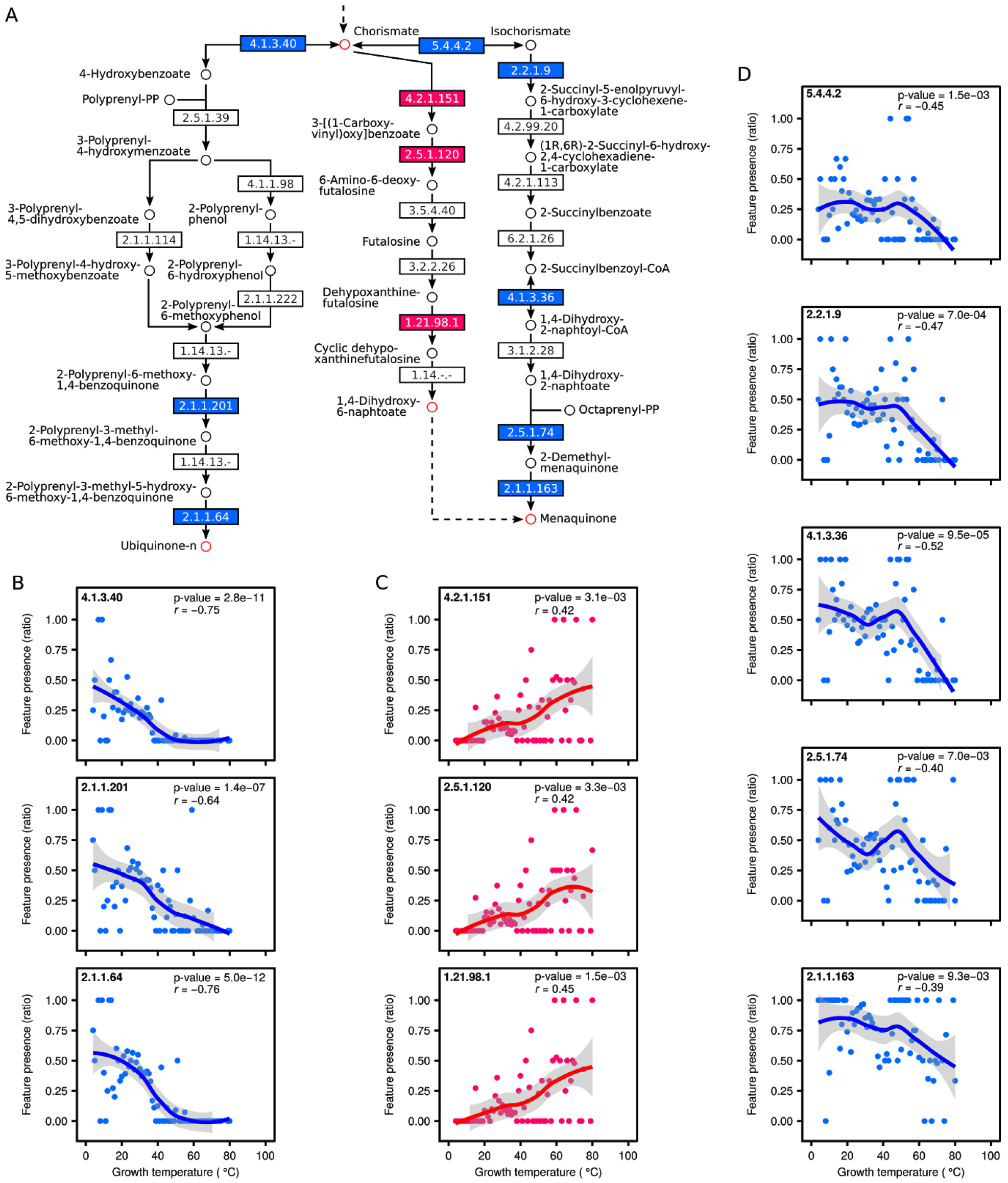
Certain metabolic pathways are enriched for the correlated enzyme activities. **(A)** Apathway diagram showing part of the KEGG pathway “ubiquinone and other terpenoid-quinone biosynthesis pathway” (map00130). Enzymes whose occurrence are significantly correlated with growth temperature (corrected p-value < 0.01) are shown in red (positive correlation) or blue (negative correlation). **(B)** The presence of enzymes participating in the biosynthesis of Ubiquinone changes with temperature. Each point indicates the occurrence of the EC number as a ratio of its presence in all organisms growing at a specific temperature. The line represents locally weighted polynomial regression with a 95% confidence band for the regression line indicated in gray. *r* indicates Spearman’s correlation coefficient. **(C)** The presence of enzymes participating in the biosynthesis of menaquinone via the futalosine pathway changes with temperature. **(D)** The presence of enzymes participating in the biosynthesis of menaquinone via the classical pathway changes with temperature.

### Domains of unknown function correlate with temperature

The EC correlation analysis is limited in that it can only leverage enzyme activities that have been characterized. Unknown activities are “hidden” and cannot be correlated. To address this limitation I performed a final analysis where domains of unknown function (DUFs) were correlated with temperature in the same manner as the EC numbers. DUFs are domains that show a conserved pattern in protein primary sequences, but for which the function of proteins in which they occur is not known. These are advantageous for the analysis insofar as they can be identified from sequence data alone and one is therefore not limited what is known through experimentation. Out of 3,918 total DUFs, 98 were were significantly correlated with temperature (Figure S3), 33 of which also had a wide phylogenetic distribution (Figure 5, Table S1). Four of the 33 were positively correlated in archaea, and eight were negatively correlated (Figure 5A). Eight were positively correlated in bacteria and seventeen were negatively correlated (Figure 5B). Two of these, DUF438 (PF04282) and DUF1957 (PF09210), were positively correlated in both archaea and bacteria. Also for negative correlation two domains, DUF3458_C (PF17432) and DUF1524 (PF07510), appeared in both archaea and bacteria.

**Figure 5.**
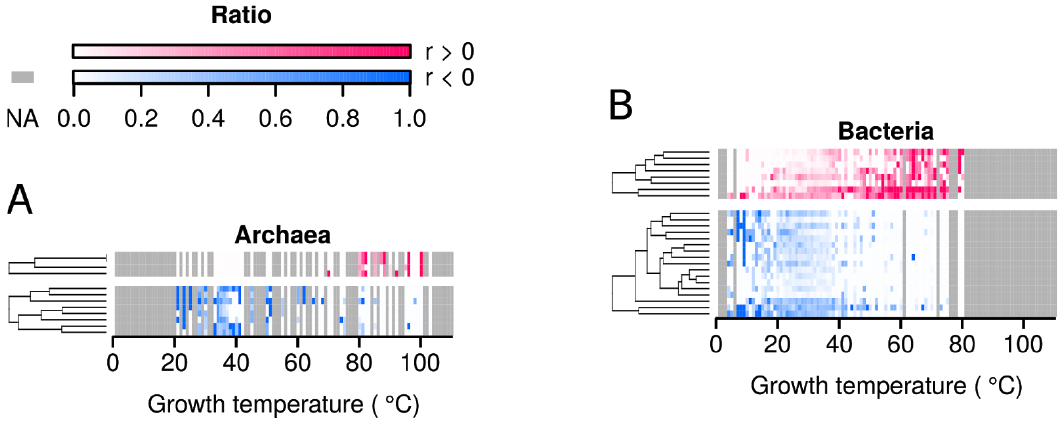
Domains of unknown function correlate with temperature. **(A)** Heat-maps showing how the occurrence of significant DUFs change with growth temperatures in archaea. Each row represents a different DUF and each column a different growth temperature. All included DUFs are statistically correlated (corrected p-value < 0.01) either positively (red) or negatively (blue) with growth temperature and a minimum phylogenetic distribution score of six (see Methods for details). For each reported growth temperature, the ratio of organisms containing each specific DUF annotation is shown by the intensity of the color. Gray indicates temperatures for which no growth temperatures are available in the dataset. **(B)** Analysis and color scale as in A, but performed with data from bacteria.

## Discussion

In this research project a dataset of 21,498 organism names and growth temperatures has been assembled by mining publicly available data from culture collection centers. This is the largest such collection of data published to date. Several other resources provide similar data, but with a more limited scope. For example; BacDive (Söhngen et al. 2014, 2015) contains growth temperatures for ~10,000 species, MediaDB (Richards et al. 2014) for less than 200 species and finally, the IJSEM phenotypic database provide the growth temperature for 4,365 strains, with the number of represented species being fewer (Barberán et al. 2017). There is a very high proportion of mesophilic organisms compared to thermophilic ones in the collected dataset (Figure 1C). The sharp drop in growth temperatures above 40 °C closely matches that seen by Barberan and collegues (Barberán et al. 2017). To what extent this reflects a true distribution of growth temperatures of naturally occurring organisms, and to what extent this is a bias introduced by what researchers have chosen to study is unclear.

For individual enzymes I found many with optima strongly differing from the growth temperature, regardless of the growth temperature of the organism from which they come (Figure 2D). The trend for mesophilic proteins to be catalytically active at higher temperatures than expected matches the observation made by Dehouck and colleagues (Dehouck, Folch, and Rooman 2008). Conversely, enzymes from thermophiles and hyperthermophiles may rely on external factors for their stability - in addition to adaptations in protein sequence and fold. This could include the action of compatible solutes (da Costa, Santos, and Galinski 1998) such as diglycerol phosphate (Lamosa et al. 2000), di-myo-inositol-phosphate (Martins et al. 1996), through increased action by chaperones, or higher protein turnover rates. Together, the unexpectedly high stability of many enzymes coming from mesophilic organisms, and the unexpectedly low stability of many of the enzymes from thermophilic and hyperthermophilic organisms likely explains the smoothing above 40 °C of the enzyme temperature histogram (Figure 2B), as compared to the growth temperature histogram (Figure 2A).

The weak correlation between single enzyme optima and growth temperature has been shown previously, albeit with a smaller dataset (Dehouck, Folch, and Rooman 2008). However, a novel insight gained in this work is that the average catalytic optimum of at least five enzymes correlates strongly with growth temperature throughout the organisms analyzed here (Figure 2E and F). This provides an important validation of the accuracy of the growth temperature dataset.

For both archaea and bacteria there are several times more enzymes that are negatively correlated with temperature (more present at low growth temperatures) than there are positively correlated ones (Figure 3C and D; Supplemental file S1). Specifically, there were 64 enzyme activities that were positively correlated with temperature and 276 that were negatively correlated. I speculate that this difference is a direct result of mesophiles having been studied to a higher degree than thermophiles and hyperthermophiles, resulting in a greater number of known enzyme activities important for growth at mesophilic temperatures. In a logical extension of this argument I propose that a large number of enzyme activities likely remain to be discovered in thermophilic organisms. Each of the 319 enzymes here shown to be differentially present at various growth temperatures highlight important targets for future hypothesis generation and experimental investigation. In particular, further study is needed to determine whether the correlation reflects true causality.

Of the eight KEGG pathways - with significant over-representation of enzymes changing with temperature - four were previously identified as over-represented in metagenomic studies of high temperature habitats (Badhai, Ghosh, and Das 2015; Saxena et al. 2016): carbon fixation pathway in prokaryotes, pyruvate metabolism, methane metabolism and purine metabolism. The TCA cycle, another enriched KEGG pathway, is known to be extensively utilized in three thermophilic bacteria (Cordova et al. 2017). In my approach, by identifying pathways with enzymes whose occurrence strongly correlates with growth temperatures, I strengthen the argument that these pathways are under evolutionary pressure in temperature adaptation. It therefore represents a clear advancement over comparing small numbers of mesophilic and thermophilic organisms and highlights the parts of metabolism under higher evolutionary pressure to change with temperature.

One of the enriched pathways identified here contains enzymes of quinone metabolism. A minority of bacteria synthesise ubiquinone (Collins and Jones 1981; Hiraishi 1999). The results obtained in this study show that among these that do synthesise it, the occurrence of ubiquinone biosynthesis genes decrease with temperature (Figure 4A and B). Menaquinone synthesis occurs via two pathways, classical and futalosine, and genes involved in the futalosine pathway are present in a range of thermophilic microorganisms (Hiratsuka et al. 2008). Here, I provide evidence that the futalosine pathway is not only present in thermophilic organisms, but is in fact the prevailing one at high growth temperatures in bacteria. I speculate that this may be an evolutionary adaptation reflecting the fact that the classical pathway contains the termolabile intermediate isochorismate (Fang, Langman, and Palmer 2010).

To expand my analysis to protein functions that have not been determined I correlated the occurrence of domains of unknown function (DUFs) with temperature. The 33 significant domains identified (Figure 5, Table S1) may represent functions important to adaptation to growth at diverse temperatures. Using computational approaches to gain insights regarding the function of DUFs has been done previously. For example, a list of 238 essential DUFs (eDUFS), were identified based on their presence in essential proteins in bacteria (Goodacre, Gerloff, and Uetz 2013). Another approach made use of remote similarity detection to establish structure-function relationships for 614 DUF families, thus providing clues to their function (Mudgal et al. 2015). Experimental approaches, notably that of structural genomics, has also been employed to gain additional information on DUFs (Jaroszewski et al. 2009). To my knowledge the approach outlined here is the first to provide evidence connecting DUFs with putative temperature adaptations in bacteria and archaea. As such, these 33 domains provide important starting points for future studies.

In this study I collected a growth temperature dataset and used it to highlight metabolic- and domain-level adaptations to growth at different temperatures. I believe that the dataset will find additional, important, uses in correlating genomic, transcriptomic, proteomic, metabolomic, phenotypic or taxonomic properties with temperature in future studies.

## Methods

### Obtaining the growth temperature dataset

Organism growth condition data was downloaded from the four culture collection centers ATCC, DSMZ, NCTC and NIES in the form of html files. For each organism record the scientific name and growth temperature was extracted using custom scripts in Python. Data relating to organism names and growth temperatures from the Pasteur institute was obtained as an Excel file. Data from the BacDive database was obtained using custom scripts in Python by interfacing with the public application programming interface (API). Records from ATCC, DSMZ and BacDive reflect those available in July of 2017. Records from Pasteur, NCTC and NIES reflect those available in the first months of 2015. Subspecies names or strain designations were removed from all organisms and the reported growth temperatures for records with the same species name were averaged and rounded to the closest integer. Organism names were further matched to taxonomic identifiers (TaxId) using the NCBI taxonomic database from July 2017; organisms for which none could be obtained were removed from the dataset. The taxonomic lineage for each organism was subsequently obtained by querying the NCBI database with the TaxIds and using the eutils resource (https://www.ncbi.nlm.nih.gov/). All growth temperatures were validated to ensure that they fall in the range of −5 to 130 °C.

### Obtaining the enzyme temperature dataset

All available experimentally determined enzyme temperature optima were extracted from the BRENDA enzyme database release 2017.2 (July 2017) with Python scripts using the Zolera SOAP package (https://pypi.python.org/pypi/ZSI/) interacting with the public BRENDA API. To de-duplicate data coming from the same enzyme the temperature optima from enzymes with the same EC number were averaged within each organism and rounded to the closest integer. TaxId was obtained for each organism using the NCBI eutils resource. Organisms for which no TaxId could be found were removed from the dataset.

### Comparing the growth- and enzyme temperature datasets

For each organism the collected growth temperatures were compared with enzyme temperature optima. The organisms present in both datasets were identified through simple matching of species names. A simple distance score was computed by subtracting each organisms growth temperature from each individual enzymes temperature optimum. Positive distance scores represent enzymes that have catalytic optima higher than the growth temperature. Negative scores represent enzymes that have optima lower than the organism growth temperature.

To compare the mean enzyme temperature optima with growth temperatures the following steps were followed: For each of the 1,811 organisms present in both the growth temperature and enzyme temperature datasets n enzyme optima, for each n is in the range 1 to 100, enzymes were sampled at random without replacement and the mean temperature calculated. In each iteration any organism having fewer than n reported enzyme optima were excluded from the calculations. The Pearson correlation between these mean enzyme temperature optima and the organism growth temperatures were subsequently calculated. The calculation was repeated 1,000 times for each n and the mean of these calculations is reported.

The mean enzyme temperature optima used in Figure 2F were calculated through simple averaging of all reported enzyme optima from each organism. Organisms having fewer than five enzyme optima were excluded. A locally weighted polynomial regression (LOESS) was calculated using the loess() function in R.

### Obtaining and filtering UniProt data

SwissProt and TrEMBL protein sequences were downloaded as fasta files from UniProt release 2017_07. A series of data matching and filtering steps were performed, as outlined below, to obtain a set of EC numbers and Pfam domains to analyze. The fasta files from both resources were parsed to extract the species name belonging to each UniProt sequence identifier. Where possible, each of these species were assigned a growth temperature through name matching with the growth temperature dataset. UniProt identifiers belonging to organisms with no assigned growth temperature were removed from further analysis. Additionally, identifiers belonging to organisms with fewer than 1,000 sequences in the resource were discarded. For the remaining UniProt identifiers EC annotations and Pfam domains, where available, were obtained using the UniProt ID mapping tool (https://www.uniprot.org/uploadlists/). For the EC dataset UniProt identifiers not annotated with an EC number and those with incomplete EC numbers (for example 1.14.-.-) were discarded. For the Pfam domain dataset UniProt identifiers not annotated with a Pfam domain were discarded. The remaining UniProt identifiers, all annotated with a species name, growth temperature, and either an EC number or Pfam domain were each subjected to correlation analysis with growth temperature.

### EC correlation analysis

The filtered UniProt EC dataset described above is skewed, disproportionately containing identifiers annotated with growth temperatures in the range 20 to 40 °C. To remove this skewing the ratio between the number of organisms carrying a protein with a specific EC annotation versus the number of organisms that do not was calculated - separately at each growth temperature. This results in a dataset where the occurrence of each EC number is represented by a single ratio, between 0.0 and 1.0, at each growth temperature. The Spearman correlation between this ratio and the growth temperature was subsequently calculated for each EC number. In this analysis identifiers from bacteria and archaea were treated separately. This was done since the growth temperature distribution of archaea and bacteria differ, with more archaea growing high temperatures and more bacteria growing at low temperatures. Analyzing the identifiers from these domains together may thus result in the identification of EC numbers that differ between bacteria and archaea. The resulting p-values were corrected for multiple testing using false detection rate (FDR). This analysis ultimately generates a set of EC numbers either significantly positively or negatively correlated with temperature. In a final step, EC numbers with a narrow phylogenetic distribution were removed. Organism relatedness was calculated from the KEGG taxonomy (http://www.genome.jp/kegg/genome.html). Organisms were treated as leaves in a rooted tree. A phylogenetic distance score that represents the amount of nodes that two leaves (organisms) are separated by was calculated. The distance will be zero for two organisms that share the same taxonomic levels but just differ on the organism name level. The maximum score in this scheme is eight. The significant EC numbers were filtered such that only those present in at least two organisms with a maximum distance of six or more were retained.

### Domain correlation analysis

The filtered UniProt Pfam domain dataset described above was filtered to retain only those corresponding to DUFs. The ratio between the number of organisms carrying a protein with a specific domain annotation versus the number of organisms that do not was calculated as described above. Calculation of the correlation of these ratios with temperature, p-value correction, and filtering for phylogenetic diversity was likewise performed as described above.

### Identifying metabolic pathways with significantly over-represented temperature-correlated enzyme activities

To identify metabolic pathways, or collection of such pathways, with statistically significant over-representation of temperature-correlated EC numbers the following steps were undertaken: All available KEGG pathway data was downloaded as xml files using the KEGG representational state transfer (REST) API (http://www.kegg.jp/kegg/rest/keggapi.html). The EC numbers listed in each of these pathways were extracted using a Python script. The occurrence of EC numbers in each of these pathways were matched against the EC numbers that significantly correlate with temperature to find the overlap. Finally, a Fischer hypergeometric test was performed, using the phyper() function from the stats package in R, to test for enrichment of temperature-correlated EC numbers in these pathways. The resulting p-values were adjusted for multiple testing using FDR and those pathways with a p-value smaller than 0.05 reported.

## Acknowledgements

I would like to thank Elzbieta Rembeza for her extensive feedback during the preparation of the manuscript, Kersten Rabe and Aleksej Zelezniak for constructive discussions and feedback during this project. I would like to thank Jens Nielsen for supporting this project with advice, time and resources. Finally, I thank Iván Domenzain for sharing the KEGG organism distance matrix used in this work.

## Supplementary figures

**Figure S1.**
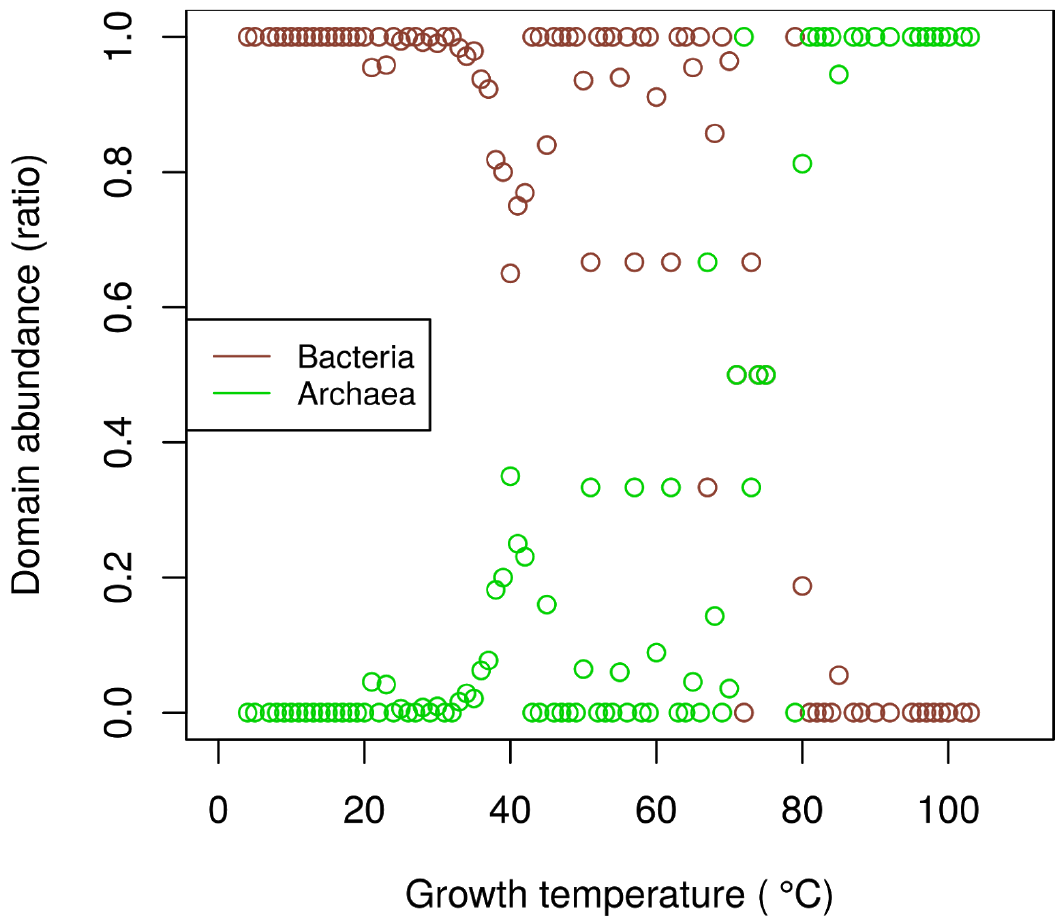
The abundance of archaeal species compared to bacterial species changes with temperature. Each point indicates the ratio of bacterial and archaeal species as a proportion of the total number of species for each of the growth temperatures in the dataset.

**Figure S2.**
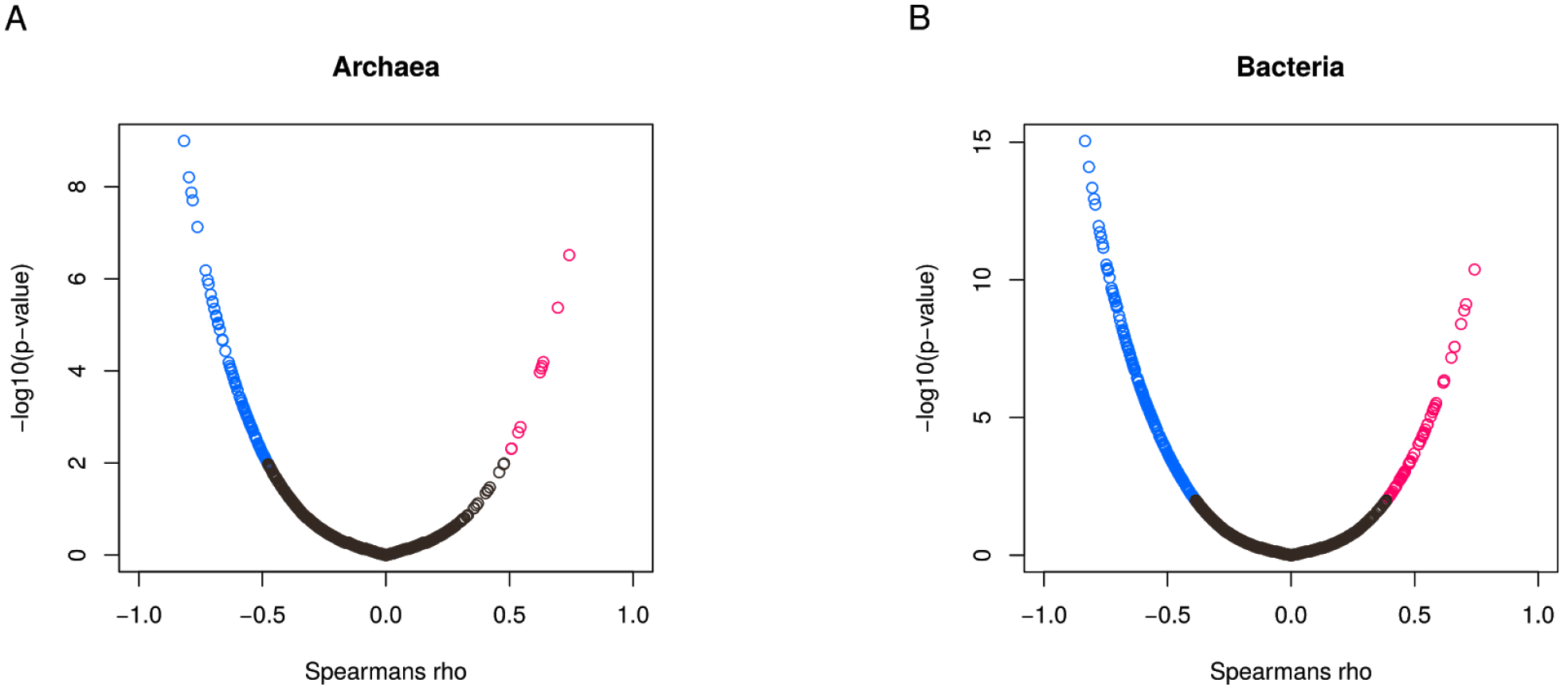
EC numbers both negatively and positively correlated with growth temperature can be identified. **(A)** The correlation between the occurrence of unique EC number annotations in species and their growth temperature is shown in archaea. Each point indicates the Spearman correlation coefficient and the corrected p-value (adjusted by false discovery rate) for a single EC number. Significant EC numbers (corrected p-value < 0.01) with positive correlation are colored red, those with negative correlation are colored blue. **(B)** The correlation between the occurrence of unique EC number annotations in species and their growth temperature is shown in bacteria Analysis and color scale as in B.

**Figure S3.**
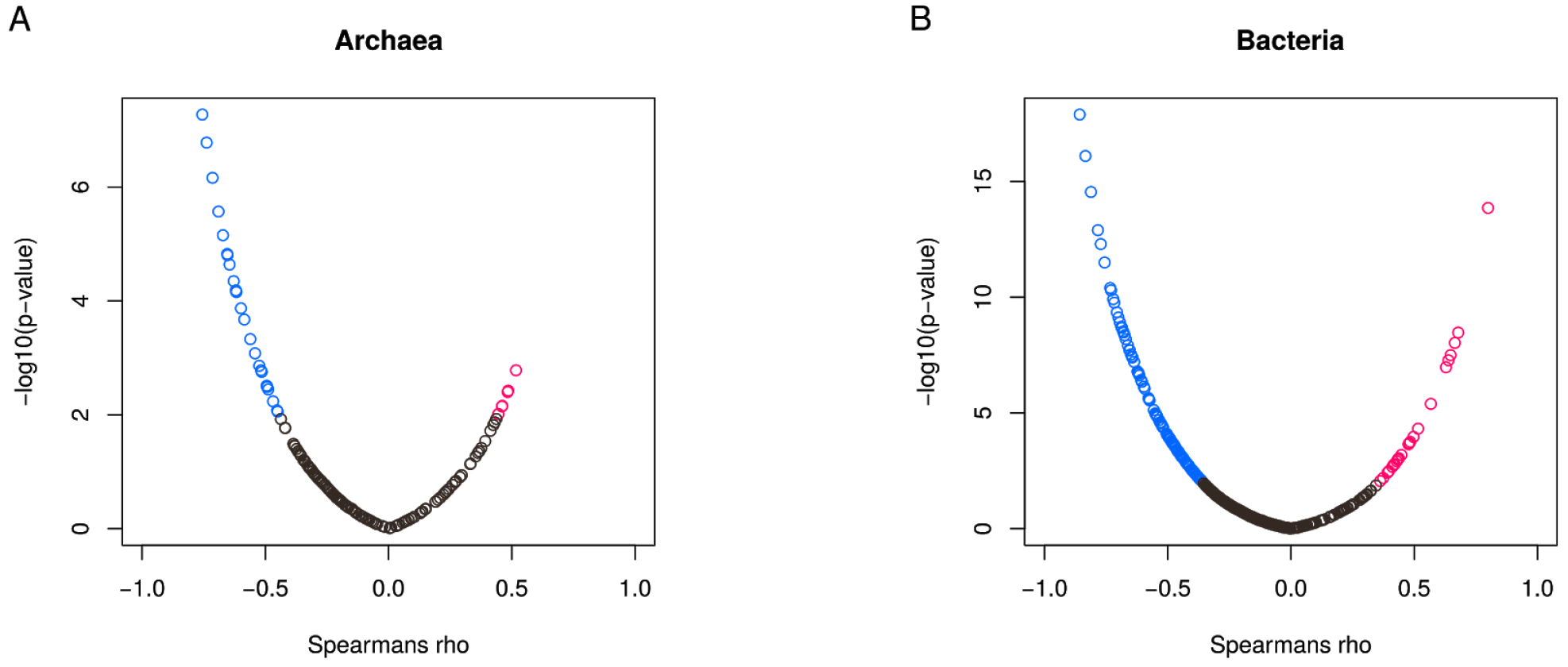
Domains of unknown function (DUFs) both negatively and positively correlated with growth temperature can be identified. **(A)** The correlation between the occurrence of unique DUFs in species and their growth temperature is shown in archaea. Each point indicates the Spearman correlation coefficient and the corrected p-value (adjusted by false discovery rate) for a single EC number. Significant EC numbers (corrected p-value < 0.01) with positive correlation are colored red, those with negative correlation are colored blue. **(B)** The correlation between the occurrence of unique DUFs in species and their growth temperature is shown in bacteria Analysis and color scale as in B.

**Table S1.**
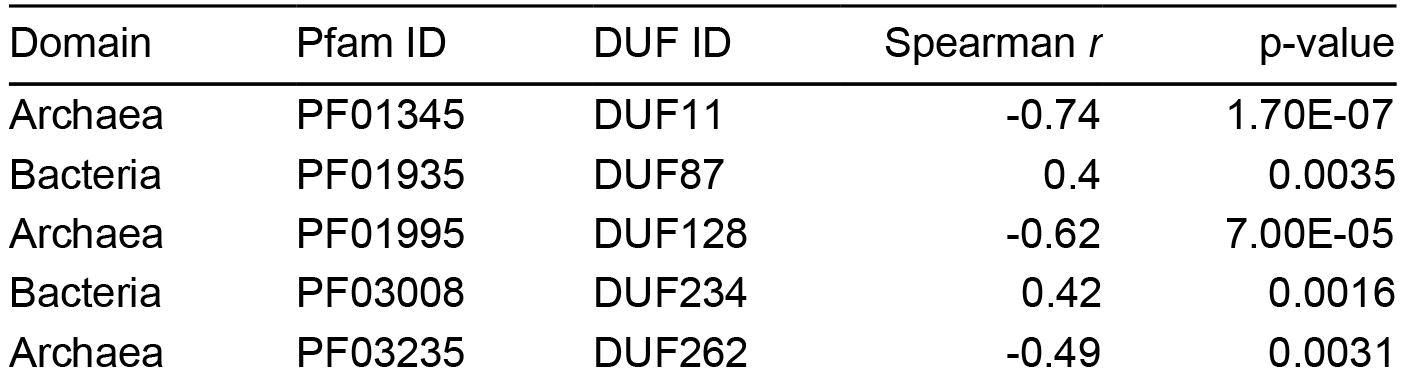
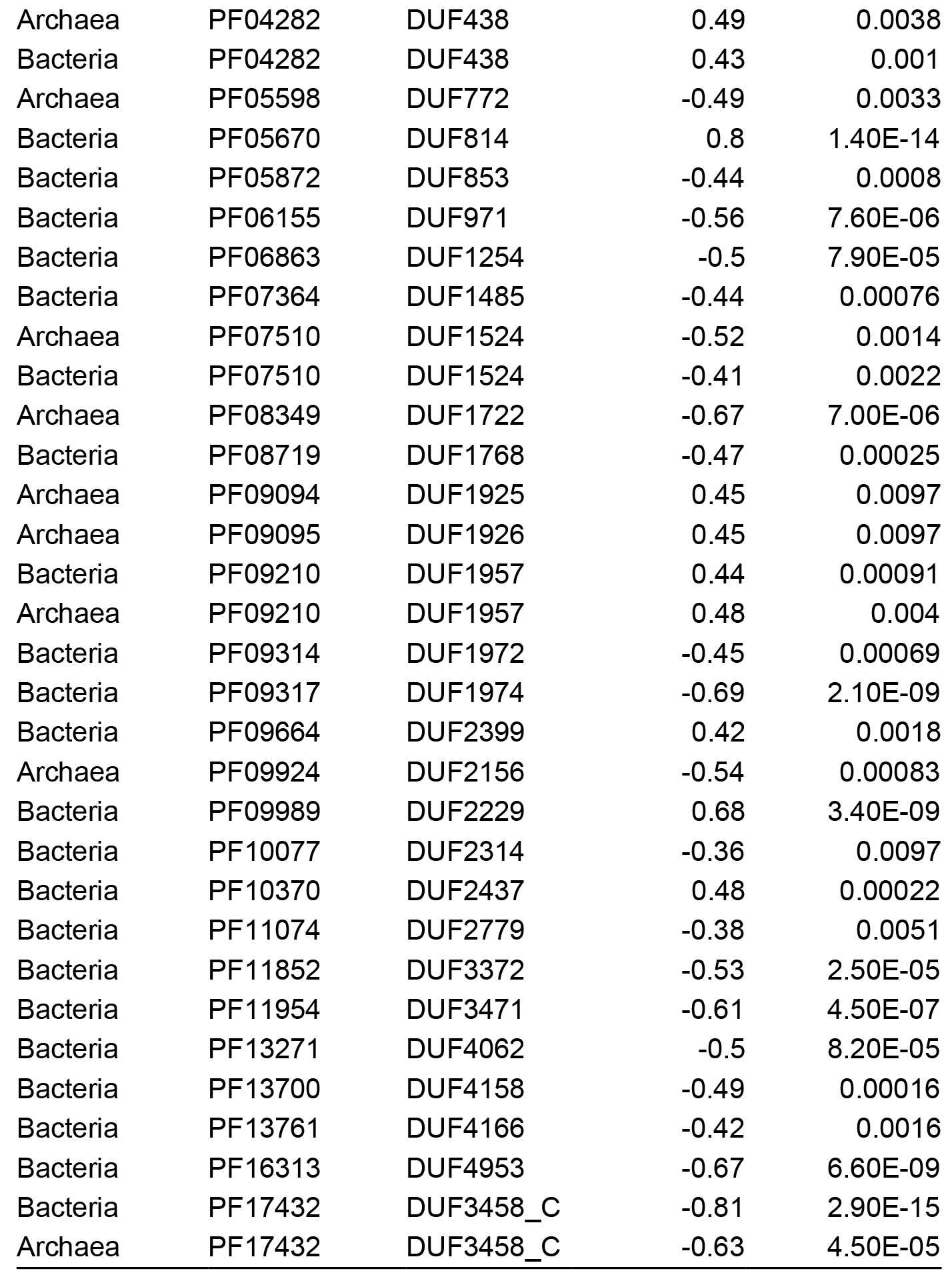
List of significant DUFs. A total of 33 unique DUFs are significantly correlated with temperature in archaea or bacteria. The table indicates the Spearman correlation coefficient and the corrected p-value (adjusted by false discovery rate). Only DUFs with a p-value less than or equal to 0.01 are included.

| Domain | Pfam ID | DUF ID | Spearman *r* | p-value |
| --- | --- | --- | --- | --- |
| Archaea | PF01345 | DUF11 | -0.74 | 1.70E-07 |
| Bacteria | PF01935 | DUF87 | 0.4 | 0.0035 |
| Archaea | PF01995 | DUF128 | -0.62 | 7.00E-05 |
| Bacteria | PF03008 | DUF234 | 0.42 | 0.0016 |
| Archaea | PF03235 | DUF262 | -0.49 | 0.0031 |
| Archaea | PF04282 | DUF438 | 0.49 | 0.0038 |
| Bacteria | PF04282 | DUF438 | 0.43 | 0.001 |
| Archaea | PF05598 | DUF772 | -0.49 | 0.0033 |
| Bacteria | PF05670 | DUF814 | 0.8 | 1.40E-14 |
| Bacteria | PF05872 | DUF853 | -0.44 | 0.0008 |
| Bacteria | PF06155 | DUF971 | -0.56 | 7.60E-06 |
| Bacteria | PF06863 | DUF1254 | -0.5 | 7.90E-05 |
| Bacteria | PF07364 | DUF1485 | -0.44 | 0.00076 |
| Archaea | PF07510 | DUF1524 | -0.52 | 0.0014 |
| Bacteria | PF07510 | DUF1524 | -0.41 | 0.0022 |
| Archaea | PF08349 | DUF1722 | -0.67 | 7.00E-06 |
| Bacteria | PF08719 | DUF1768 | -0.47 | 0.00025 |
| Archaea | PF09094 | DUF1925 | 0.45 | 0.0097 |
| Archaea | PF09095 | DUF1926 | 0.45 | 0.0097 |
| Bacteria | PF09210 | DUF1957 | 0.44 | 0.00091 |
| Archaea | PF09210 | DUF1957 | 0.48 | 0.004 |
| Bacteria | PF09314 | DUF1972 | -0.45 | 0.00069 |
| Bacteria | PF09317 | DUF1974 | -0.69 | 2.10E-09 |
| Bacteria | PF09664 | DUF2399 | 0.42 | 0.0018 |
| Archaea | PF09924 | DUF2156 | -0.54 | 0.00083 |
| Bacteria | PF09989 | DUF2229 | 0.68 | 3.40E-09 |
| Bacteria | PF10077 | DUF2314 | -0.36 | 0.0097 |
| Bacteria | PF10370 | DUF2437 | 0.48 | 0.00022 |
| Bacteria | PF11074 | DUF2779 | -0.38 | 0.0051 |
| Bacteria | PF11852 | DUF3372 | -0.53 | 2.50E-05 |
| Bacteria | PF11954 | DUF3471 | -0.61 | 4.50E-07 |
| Bacteria | PF13271 | DUF4062 | -0.5 | 8.20E-05 |
| Bacteria | PF13700 | DUF4158 | -0.49 | 0.00016 |
| Bacteria | PF13761 | DUF4166 | -0.42 | 0.0016 |
| Bacteria | PF16313 | DUF4953 | -0.67 | 6.60E-09 |
| Bacteria | PF17432 | DUF3458_C | -0.81 | 2.90E-15 |
| Archaea | PF17432 | DUF3458_C | -0.63 | 4.50E-05 |

## References

Aravind, L., R. L. Tatusov, Y. I. Wolf, D. R. Walker, and E. V. Koonin. 1998. “Evidence for Massive Gene Exchange between Archaeal and Bacterial Hyperthermophiles.” Trends in Genetics: TIG 14 (11):442–44.

Badhai, Jhasketan, Tarini S. Ghosh, and Subrata K. Das. 2015. “Taxonomic and Functional Characteristics of Microbial Communities and Their Correlation with Physicochemical Properties of Four Geothermal Springs in Odisha, India.” Frontiers in Microbiology 6:1166.

Barberán, Albert, Hildamarie Caceres Velazquez, Stuart Jones, and Noah Fierer. 2017. “Hiding in Plain Sight: Mining Bacterial Species Records for Phenotypic Trait Information.” mSphere 2 (4). https://doi.org/10.1128/mSphere.00237-17.

Bezsudnova, Ekaterina Y., Konstantin M. Boyko, Konstantin M. Polyakov, Pavel V. Dorovatovskiy, Tatiana N. Stekhanova, Vadim M. Gumerov, Nikolai V. Ravin, Konstantin G. Skryabin, Michael V. Kovalchuk, and Vladimir O. Popov. 2012. “Structural Insight into the Molecular Basis of Polyextremophilicity of Short-Chain Alcohol Dehydrogenase from the Hyperthermophilic Archaeon Thermococcus Sibiricus.” Biochimie 94 (12):2628–38.

Boutz, Daniel R., Duilio Cascio, Julian Whitelegge, L. Jeanne Perry, and Todd O. Yeates. 2007. “Discovery of a Thermophilic Protein Complex Stabilized by Topologically Interlinked Chains.” Journal of Molecular Biology 368 (5):1332–44.

Brasen, C., D. Esser, B. Rauch, and B. Siebers. 2014. “Carbohydrate Metabolism in Archaea: Current Insights into Unusual Enzymes and Pathways and Their Regulation.” Microbiology and Molecular Biology Reviews: MMBR 78 (1):89–175.

Brbić, Maria, Matija Piškorec, Vedrana Vidulin, Anita Kriško, Tomislav Šmuc, and Fran Supek. 2016. “The Landscape of Microbial Phenotypic Traits and Associated Genes.” Nucleic Acids Research 44 (21):10074–90.

Brumm, Phillip J., Scott Monsma, Brendan Keough, Svetlana Jasinovica, Erin Ferguson, Thomas Schoenfeld, Michael Lodes, and David A. Mead. 2015. “Complete Genome Sequence of Thermus Aquaticus Y51MC23.” PloS One 10 (10):e0138674.

Burra, Prasad V., Lajos Kalmar, and Peter Tompa. 2010. “Reduction in Structural Disorder and Functional Complexity in the Thermal Adaptation of Prokaryotes.” PloS One 5 (8):e12069.

Collins, M. D., and D. Jones. 1981. “Distribution of Isoprenoid Quinone Structural Types in Bacteria and Their Taxonomic Implication.” Microbiological Reviews 45 (2):316–54.

Cordova, Lauren T., Robert M. Cipolla, Adti Swarup, Christopher P. Long, and Maciek R. Antoniewicz. 2017. “13C Metabolic Flux Analysis of Three Divergent Extremely Thermophilic Bacteria: Geobacillus Sp. LC300, Thermus Thermophilus HB8, and Rhodothermus Marinus DSM 4252.” Metabolic Engineering 44 (November):182–90.

Costa, M. S. da, H. Santos, and E. A. Galinski. 1998. “An Overview of the Role and Diversity of Compatible Solutes in Bacteria and Archaea.” Advances in Biochemical Engineering/biotechnology 61: 117–53.

Counts, James A., Benjamin M. Zeldes, Laura L. Lee, Christopher T. Straub, Michael W. W. Adams, and Robert M. Kelly. 2017. “Physiological, Metabolic and Biotechnological Features of Extremely Thermophilic Microorganisms.” Wiley Interdisciplinary Reviews. Systems Biology and Medicine 9 (3). https://doi.org/10.1002/wsbm.1377.

Cowan, D. A., J-B Ramond, T. P. Makhalanyane, and P. De Maayer. 2015. “Metagenomics of Extreme Environments.” Current Opinion in Microbiology 25 (June):97–102.

DeCastro, María-Eugenia, Esther Rodríguez-Belmonte, and María-Isabel González-Siso. 2016. “Metagenomics of Thermophiles with a Focus on Discovery of Novel Thermozymes.” Frontiers in Microbiology 7 (September):1521.

Dehouck, Yves, Benjamin Folch, and Marianne Rooman. 2008. “Revisiting the Correlation between Proteins’ Thermoresistance and Organisms’ Thermophilicity.” Protein Engineering, Design & Selection: PEDS 21 (4):275–78.

Ettema, Thijs J. G., Hatim Ahmed, Ans C. M. Geerling, John van der Oost, and Bettina Siebers. 2008. “The Non-Phosphorylating Glyceraldehyde-3-Phosphate Dehydrogenase (GAPN) of Sulfolobus Solfataricus: A Key-Enzyme of the Semi-Phosphorylative Branch of the Entner-Doudoroff Pathway.” Extremophiles: Life under Extreme Conditions 12 (1):75–88.

Fang, Maohai, Blaine M. Langman, and David R. J. Palmer. 2010. “A Stable Analog of Isochorismate for the Study of MenD and Other Isochorismate-Utilizing Enzymes.” Bioorganic & Medicinal Chemistry Letters 20 (17):5019–22.

Forterre, P., C. Bouthier De La Tour, H. Philippe, and M. Duguet. 2000. “Reverse Gyrase from Hyperthermophiles: Probable Transfer of a Thermoadaptation Trait from Archaea to Bacteria.” Trends in Genetics: TIG 16 (4):152–54.

Goodacre, N. F., D. L. Gerloff, and P. Uetz. 2013. “Protein Domains of Unknown Function Are Essential in Bacteria.” mBio 5 (1):e00744–13 – e00744–13.

Henne, Anke, Holger Brüggemann, Carsten Raasch, Arnim Wiezer, Thomas Hartsch, Heiko Liesegang, Andre Johann, et al. 2004. “The Genome Sequence of the Extreme Thermophile Thermus Thermophilus.” Nature Biotechnology 22 (April). Nature Publishing Group:547.

Hiraishi, A. 1999. “Isoprenoid Quinones as Biomarkers of Microbial Populations in the Environment.” Journal of Bioscience and Bioengineering 88 (5):449–60.

Hiratsuka, Tomoshige, Kazuo Furihata, Jun Ishikawa, Haruyuki Yamashita, Nobuya Itoh, Haruo Seto, and Tohru Dairi. 2008. “An Alternative Menaquinone Biosynthetic Pathway Operating in Microorganisms.” Science 321 (5896):1670–73.

Jaroszewski, Lukasz, Zhanwen Li, S. Sri Krishna, Constantina Bakolitsa, John Wooley, Ashley M. Deacon, Ian A. Wilson, and Adam Godzik. 2009. “Exploration of Uncharted Regions of the Protein Universe.” PLoS Biology 7 (9):e1000205.

Kampmann, Martin, and Daniela Stock. 2004. “Reverse Gyrase Has Heat-Protective DNA Chaperone Activity Independent of Supercoiling.” Nucleic Acids Research 32 (12):3537–45.

Kanehisa, Minoru, Miho Furumichi, Mao Tanabe, Yoko Sato, and Kanae Morishima. 2017. “KEGG: New Perspectives on Genomes, Pathways, Diseases and Drugs.” Nucleic Acids Research 45 (D1):D353–61.

Kengen, S. W., F. A. de Bok, N. D. van Loo, C. Dijkema, A. J. Stams, and W. M. de Vos. 1994. “Evidence for the Operation of a Novel Embden-Meyerhof Pathway That Involves ADP-Dependent Kinases during Sugar Fermentation by Pyrococcus Furiosus.” The Journal of Biological Chemistry 269 (26):17537–41.

Koga, Yosuke. 2012. “Thermal Adaptation of the Archaeal and Bacterial Lipid Membranes.” Archaea 2012 (August):789652.

Kosuge, T., and T. Hoshino. 1998. “Lysine Is Synthesized through the Alpha-Aminoadipate Pathway in Thermus Thermophilus.” FEMS Microbiology Letters 169 (2):361–67.

Lamosa, P., A. Burke, R. Peist, R. Huber, M. Y. Liu, G. Silva, C. Rodrigues-Pousada, J. LeGall, C. Maycock, and H. Santos. 2000. “Thermostabilization of Proteins by Diglycerol Phosphate, a New Compatible Solute from the Hyperthermophile Archaeoglobus Fulgidus.” Applied and Environmental Microbiology 66 (5):1974–79.

Lee, Na-Rae, Meiyappan Lakshmanan, Shilpi Aggarwal, Ji-Won Song, Iftekhar A. Karimi, Dong-Yup Lee, and Jin-Byung Park. 2014. “Genome-Scale Metabolic Network Reconstruction and in Silico Flux Analysis of the Thermophilic Bacterium Thermus Thermophilus HB27.” Microbial Cell Factories 13 (April):61.

Le, Phuong Thi, Thulani P. Makhalanyane, Leandro D. Guerrero, Surendra Vikram, Yves Van de Peer, and Don A. Cowan. 2016. “Comparative Metagenomic Analysis Reveals Mechanisms for Stress Response in Hypoliths from Extreme Hyperarid Deserts.” Genome Biology and Evolution 8 (9):2737–47.

Martins, L. O., L. S. Carreto, M. S. Da Costa, and H. Santos. 1996. “New Compatible Solutes Related to Di-Myo-Inositol-Phosphate in Members of the Order Thermotogales.” Journal of Bacteriology 178 (19):5644–51.

Mehta, Ridhi, Paavan Singhal, Hardeep Singh, Dhanashree Damle, and Anil K. Sharma. 2016. “Insight into Thermophiles and Their Wide-Spectrum Applications.” 3 Biotech 6 (1):81.

Mudgal, Richa, Sankaran Sandhya, Nagasuma Chandra, and Narayanaswamy Srinivasan. 2015. “De-DUFing the DUFs: Deciphering Distant Evolutionary Relationships of Domains of Unknown Function Using Sensitive Homology Detection Methods.” Biology Direct 10 (July):38.

Noort, Vera van, Bettina Bradatsch, Manimozhiyan Arumugam, Stefan Amlacher, Gert Bange, Chris Creevey, Sebastian Falk, et al. 2013. “Consistent Mutational Paths Predict Eukaryotic Thermostability.” BMC Evolutionary Biology 13 (January):7.

Ogata, H., S. Goto, K. Sato, W. Fujibuchi, H. Bono, and M. Kanehisa. 1999. “KEGG: Kyoto Encyclopedia of Genes and Genomes.” Nucleic Acids Research 27 (1):29–34.

Oshima, T. 2007. “Unique Polyamines Produced by an Extreme Thermophile, Thermus Thermophilus.” Amino Acids 33 (2):367–72.

Reimer, Lorenz C., Carola Söhngen, Anna Vetcininova, and Jörg Overmann. 2017. “Mobilization and Integration of Bacterial Phenotypic Data-Enabling next Generation Biodiversity Analysis through the BacDive Metadatabase.” Journal of Biotechnology 261 (November):187–93.

Richards, Matthew A., Victor Cassen, Benjamin D. Heavner, Nassim E. Ajami, Andrea Herrmann, Evangelos Simeonidis, and Nathan D. Price. 2014. “MediaDB: A Database of Microbial Growth Conditions in Defined Media.” PloS One 9 (8):e103548.

Sabath, Niv, Evandro Ferrada, Aditya Barve, and Andreas Wagner. 2013. “Growth Temperature and Genome Size in Bacteria Are Negatively Correlated, Suggesting Genomic Streamlining during Thermal Adaptation.” Genome Biology and Evolution 5 (5):966–77.

Saxena, Rituja, Darshan B. Dhakan, Parul Mittal, Prashant Waiker, Anirban Chowdhury, Arundhuti Ghatak, and Vineet K. Sharma. 2016. “Metagenomic Analysis of Hot Springs in Central India Reveals Hydrocarbon Degrading Thermophiles and Pathways Essential for Survival in Extreme Environments.” Frontiers in Microbiology 7:2123.

Schäfers, Christian, Saskia Blank, Sigrid Wiebusch, Skander Elleuche, and Garabed Antranikian. 2017. “Complete Genome Sequence of Thermus Brockianus GE-1 Reveals Key Enzymes of Xylan/xylose Metabolism.” Standards in Genomic Sciences 12 (February):22.

Schomburg, I., L. Jeske, M. Ulbrich, S. Placzek, A. Chang, and D. Schomburg. 2017. “The BRENDA Enzyme Information System-From a Database to an Expert System.” Journal of Biotechnology 261 (November):194–206.

Selig, M., K. B. Xavier, H. Santos, and P. Schönheit. 1997. “Comparative Analysis of Embden-Meyerhof and Entner-Doudoroff Glycolytic Pathways in Hyperthermophilic Archaea and the Bacterium Thermotoga.” Archives of Microbiology 167 (4):217–32.

Söhngen, Carola, Boyke Bunk, Adam Podstawka, Dorothea Gleim, and Jörg Overmann. 2014. “BacDive--the Bacterial Diversity Metadatabase.” Nucleic Acids Research 42 (Database issue):D592–99.

Söhngen, Carola, Adam Podstawka, Boyke Bunk, Dorothea Gleim, Anna Vetcininova, Lorenz Christian Reimer, Christian Ebeling, Cezar Pendarovski, and Jörg Overmann. 2015. “BacDive - The Bacterial Diversity Metadatabase in 2016.” Nucleic Acids Research 44 (D1):D581–85.

Stetter, Karl O. 1996. “Hyperthermophilic Procaryotes.” FEMS Microbiology Reviews 18 (2-3). Blackwell Publishing Ltd149–58.

Swarup, Aditi, Jing Lu, Kathleen C. DeWoody, and Maciek R. Antoniewicz. 2014. “Metabolic Network Reconstruction, Growth Characterization and 13C-Metabolic Flux Analysis of the Extremophile Thermus Thermophilus HB8.” Metabolic Engineering 24 (July):173–80.

UniProt Consortium. 2015. “UniProt: A Hub for Protein Information.” Nucleic Acids Research 43 (Database issue):D204–12.

Wang, Quanhui, Zhen Cen, and Jingjing Zhao. 2015. “The Survival Mechanisms of Thermophiles at High Temperatures: An Angle of Omics.” Physiology 30 (2):97–106.

Wu, Dongying, Jason Raymond, Martin Wu, Sourav Chatterji, Qinghu Ren, Joel E. Graham, Donald A. Bryant, et al. 2009. “Complete Genome Sequence of the Aerobic CO-Oxidizing Thermophile Thermomicrobium Roseum.” PloS One 4 (1):e4207.

